# Ribocutter: Cas9-mediated rRNA depletion from multiplexed Ribo-seq libraries

**DOI:** 10.1101/2021.07.14.451473

**Authors:** Oscar G. Wilkins, Jernej Ule

## Abstract

RNA sequencing libraries produced from fragmented RNA, especially Ribo-seq libraries, contain high proportions of reads from abundant non-coding RNAs. Here, we describe a streamlined Cas9-based protocol for removing abundant rRNA/ncRNA contaminants from Ribo-seq (or other small RNA-seq) libraries and an easy-to-use software tool, *ribocutter*, for designing ready-to-order sgRNA templates. Following sgRNA template design, the pool of templates is *in vitro* transcribed using a 1-step commercial kit, which produces enough sgRNAs for multiple treatments. A single multiplexed sequencing library is then treated with Cas9/sgRNAs, followed by a short PCR program, which can increase the fraction of useful reads by more than 3-fold. Comparison of samples before and after depletion demonstrates that Cas9 produces minimal off-target effects and preserves key features (eg. footprint length, periodicity) of Ribo-seq libraries. The method is thus highly effective, costs <£0.50 per sample, and minimises non-specific depletion and technical variation between samples.

## Introduction

The Ribo-seq method involves digestion of RNAs with ribonucleases, leaving only small sections of mRNA protected within ribosomes intact (Ingolia et al. 2009). These small mRNA-fragments, known as ribosome footprints (RFPs), are then size selected and sequenced. In spite of the stringent size selection, the partial digestion of the ribosomal RNA (rRNA) and other abundant non-coding RNAs (ncRNAs) results in high levels of ncRNA contamination in the libraries (McGlincy and Ingolia 2017; Zinshteyn et al. 2020; Alkan et al. 2021). Due to the narrow size selection, the contamination typically consists of a small number of distinct, highly abundant sequences; this means that rRNA depletion methods that tile the entire rRNA are very prone to oversaturation, rendering them largely ineffectual (Zinshteyn et al. 2020; Alkan et al. 2021).

The most popular approach for depleting highly-abundant contaminants from ribosome profiling libraries is subtractive hybridisation with bespoke biotinylated probes that are complementary to the abundant rRNA fragments (Ingolia et al. 2012; Zinshteyn et al. 2020; Alkan et al. 2021). However, although this approach can produce good results when well optimised, it has several disadvantages. First, it requires the most abundant contaminants to be predicted in advance, but contaminants may vary significantly between experiments and samples (Alkan et al. 2021). Second, because depletion is performed before samples can be multiplexed, each sample must be treated separately, which is labour-intensive and risks introducing technical variation between samples. Third, because hybridisation is typically performed at a much lower temperature than the predicted Tm of each bait oligo, it is possible that off-target annealing of probes will lead to depletion of non-desired fragments. Fourth, additional RNA degradation may occur via non-specific RNase activity or non-enzymatic RNA hydrolysis at the elevated temperatures used during the initial denaturation (Alkan et al. 2021).

Recently, methods have been developed that deplete contaminant sequences from the sequencing library using Cas9 (Gu et al. 2016; Montefiori et al. 2017; Hardigan et al. 2019; Han et al. 2020). This approach has several advantages compared to the use of biotinylated probes. First and foremost, a single multiplexed sample can be treated after PCR amplification of the library, or even after its first round of sequencing, thus eliminating technical variation between samples and removing the need to predict contaminants in advance. Second, Cas9-mediated degradation is highly-specific and reduces the risk of off-target degradation (Wu, Kriz, and Sharp 2014). Third, because the risk of RNA degradation is eliminated, and an aliquot of the pre-treated amplified library can be held in reserve as a backup in case of technical errors during the depletion procedure, the risk of sample loss is greatly decreased.

In spite of its many potential benefits, the Cas9-mediated method has so far been used by only one ribosome profiling study, and the protocol provided by this study is time-consuming, especially for sgRNA synthesis (Han et al. 2020). Furthermore, this study did not analyse the potential biases and off-target depletion that Cas9-mediated rRNA depletion might introduce into Ribo-seq data. Such analysis is required, especially given that RNase H-mediated rRNA depletion was recently shown to be unsuitable for Ribo-seq due to the introduced biases (Zinshteyn et al. 2020).

Here, we describe Ribocutter, a rapid, efficient and very affordable protocol for the synthesis of sgRNAs and Cas9 treatment of multiplexed sequencing libraries, along with an associated software tool for sgRNA design, available on Bioconda, PyPi and Github. Importantly, we rigorously demonstrate that our protocol does not interfere with the subcodon periodicity of RFPs and has very limited off-target depletion. Overall, our protocol greatly increases the fraction of useful reads in the library, without introducing significant experimental bias, at a negligible cost of <£0.50 per sample.

## Protocol

### Required materials

- EnGen® sgRNA Synthesis Kit, S. pyogenes (NEB, E3322V)
- 20 uM Cas9 Nuclease, S. pyogenes (NEB, M0386T or M0386M)
- Ethanol
- Totalpure NGS Beads (or equivalent, eg. Beckman Ampure XP)
- Nuclease-free water
- RNase-free tubes and tips
- RNase A (10 mg/ml)
- Phusion 2x HF master mix (or similar)
- Primers for multiplexed amplification (Table 1)
- Thermocycler

**Table 1:**
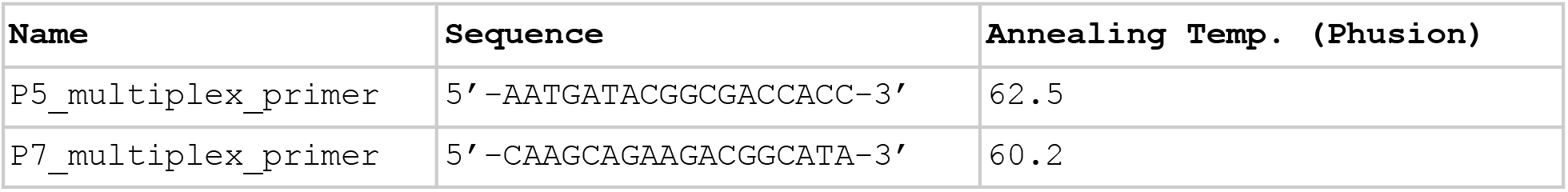
Example primers that could be used for amplification of a multiplexed sequencing library, in this case with P5 and P7 Illumina overhangs (Neckles et al. 2019) (the typical Illumina configuration).

### Optional materials

- Murine RNase Inhibitor (NEB, M0314S)

### Overview

The protocol consists of four steps. The first two steps produce sgRNAs, and thus are optional if one can use a sgRNA pool that has been effective for previous treatments of similar libraries. The subsequent steps involve Cas9-mediated cleavage of contaminant sequences, followed by PCR amplification of uncleaved sequences (**Figure 1)**.

**Figure 1:**
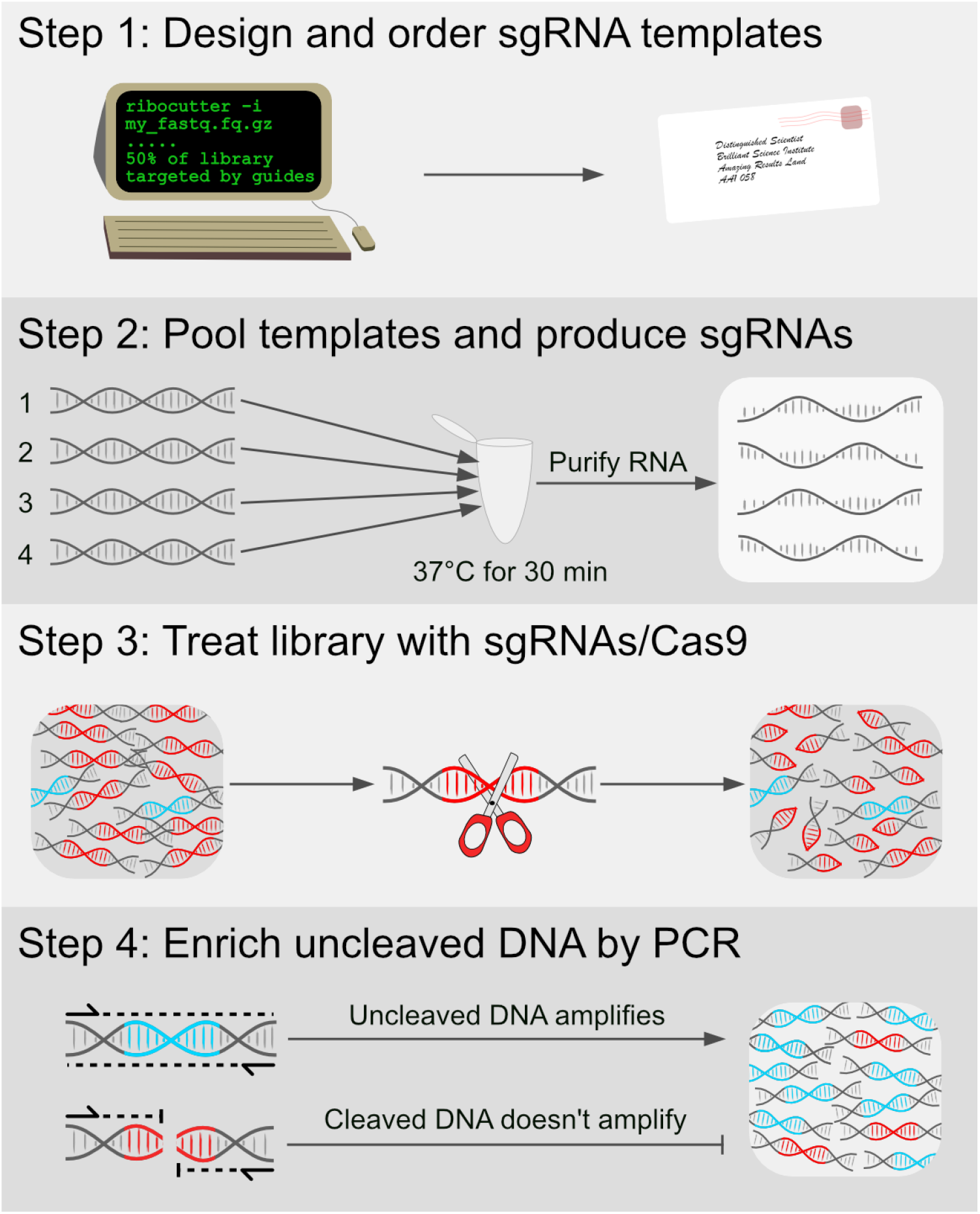
An overview of the Ribocutter method. Step 1: sgRNA templates are designed and ordered. Step 2: These are used to produce sgRNAs in a 1-step reaction. Step 3: A PCR-amplified ribosome profiling sequencing library is used as input for digestion - this library will contain a large quantity of contaminant sequences (red) and a smaller fraction of ribosome footprints (blue), both with Illumina-compatible overhangs on each end (grey). The library is treated with sgRNAs/Cas9, which cleaves the targeted contaminant sequences. Step 4: A short PCR step, using primers that anneal to the extremities of the Illumina adapters (thus preserving barcodes/indices), greatly enriches for uncleaved sequences, because cleaved amplicons cannot amplify. The final library contains a far greater proportion of ribosome footprints than the input.

### Step 1: Design of guides and purchase of template oligos

In Table S1 we have provided a list of oligos for the synthesis of sgRNAs targeting common mammalian contaminants. However, we have also made available the Ribocutter command-line tool, which automatically designs guides against the most abundant contaminants in a user-provided fastq file. The researcher can choose whether to simply order the oligos in Table S1, or use Ribocutter to determine bespoke guides for their particular samples.

1. Optional: Run *ribocutter -i adapter_trimmed_fastq.gz*, where *adapter_trimmed_fastq.gz* is a ribosome profiling sample. The fastq should have had its adapter sequences removed, for example with cutadapt (Martin 2011). Ribocutter will immediately estimate the fraction of the library that is targeted by the suggested guides; the number of guides can be specified with option -g/--guides. *Optionally, to reduce off-target effects, a fasta file of background sequences which should not be targeted can be included; however, this is not recommended*.
2. Order oligos designed by Ribocutter, using the smallest synthesis scale and cheapest purification method available; these are compatible with the EnGen sgRNA synthesis kit. *For large numbers of oligos it may be most economical to order a 50 pmol/oligo OPool from IDT*.

### Step 2: Synthesis of sgRNAs (if required*)

1. Resuspend oligos in nuclease-free water.
2. Combine oligos for a final combined concentration of 1 uM - *for example if making a pool of 10 oligos, each should be 0.1 uM on average; individual concentrations do not have to be uniform - oligos that target the most abundant contaminants may be spiked in at higher concentration*.
3. Follow the EnGen sgRNA Synthesis Kit protocol to produce sgRNAs using the 1 uM oligo pool as input; optionally include 0.5 ul of Murine RNase inhibitor to minimise the risk of sgRNA degradation.
4. Purify RNA from the DNase I-treated sample with Totalpure NGS/Ampure RNAClean beads - use 3x volume (60 ul) of beads for the 20 ul sample - *alternatively perform trizol-based purification or ethanol precipitation*
5. Resuspend in 20 ul of nuclease-free water.
6. Quantify the RNA by nanodrop (or equivalent)
7. Calculate the RNA molarity, assuming a sgRNA molecular mass of 35 kDa - *for example if 350 ng/ul, this is 10 uM*.
8. Store at −20 for short term, or −80 for long term. Aliquoting is advisable.

### Step 3: Treatment of library and removal of sgRNA and Cas9

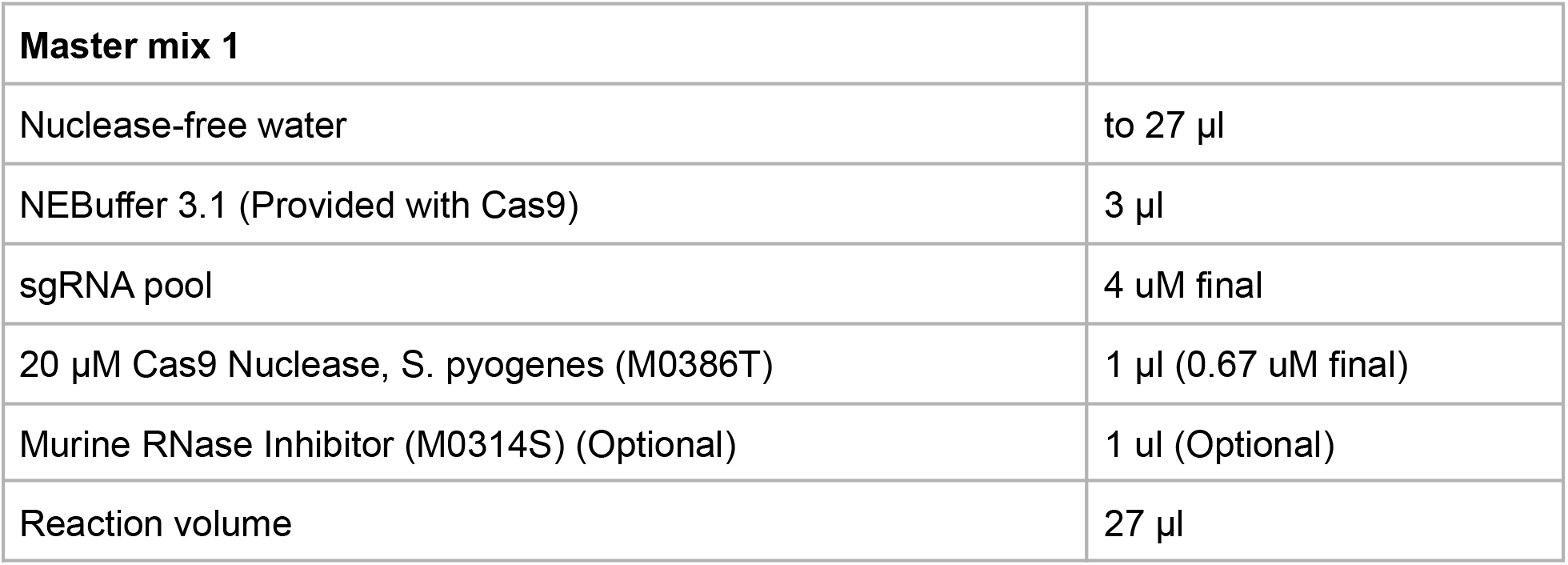

|  |  |
| --- | --- |
| **Master mix 1** |  |
| Nuclease-free water | to 27 $\mu\text{l}$ |
| NEBuffer 3.1 (Provided with Cas9) | 3 $\mu\text{l}$ |
| sgRNA pool | 4 $\mu\text{M}$ final |
| 20 $\mu\text{M}$ Cas9 Nuclease, *S. pyogenes* (M0386T) | 1 $\mu\text{l}$ (0.67 $\mu\text{M}$ final) |
| Murine RNase Inhibitor (M0314S) (Optional) | 1 $\mu\text{l}$ (Optional) |
| Reaction volume | 27 $\mu\text{l}$ |

1. Make the above master mix in the order listed
2. Pre-incubate for 10 min at 25°C
3. Add 3 ul of purified, multiplexed sequencing library (initial concentration of library 10-50 nM)
4. Incubate at 37 degrees for 30 minutes (or up to 4 h)
5. Heat inactivate Cas9 at 65 degrees for 5 min
6. Add 2 ul of RNase A (10 mg/ml) and incubate at 37 for 10 min.
7. Purify with 54 ul of Totalpure NGS/Ampure XP beads and resuspend in 10 ul of water - *alternatively perform ethanol precipitation or column-based PCR cleanup*

### Step 4: Re-amplification of the treated library**

Cleaved dsDNA will not amplify. Thus, dsDNA containing uncleaved (i.e. ribosome footprint) inserts can be enriched by PCR amplification of the treated library, using suitable primers for multiplexed PCR.

1. Estimate the number of cycles needed to produce sufficient input for Illumina sequencing, accounting for dilutions and assuming the Cas9 treatment will have destroyed 50-75% of the library. *Typically 5-7 cycles will be sufficient*.
2. Perform a high-fidelity PCR reaction with Illumina-compatible multiplex primers using the purified Cas9-treated library as input **(Table 1)**. *Because the primers anneal to the extremities of the Illumina adapters, i5/i7 indices are preserved. Alternative high-fidelity PCR mixes may be used (eg Q5)*.
3. Purify the PCR product with 1.8x Totalpure NGS/Ampure XP beads and resuspend in a suitable volume
4. Assess the concentration by Qubit/Tapestation/Quantifluor and submit for sequencing

## Results

### Cas9 enables robust and specific rRNA depletion

In our initial attempts at Cas9 depletion of rRNA/contaminant sequences, we used the concentrations of reagents and substrate for *in vitro* Cas9 reactions recommended by NEB (10:10:1 ratio of sgRNA:Cas9:target). However, while we did achieve substantial depletion of most targeted sequences, we found some sequences largely escaped degradation (data not shown). We noted that the original Cas9 targeted depletion protocol used far higher concentrations of Cas9 and sgRNAs (1000:100:1 sgRNA:Cas9:target), and thus increased the concentration of sgRNAs and Cas9 enzyme in our own reactions (Gu et al. 2016).

Using our improved, high-concentration protocol, we depleted abundant sequences from five ribosome profiling libraries derived from mouse brain. We chose these libraries because we had previously sequenced them before depletion, and because they were produced with a wide (30-fold) range of RNase concentrations that are expected to generate a variety of abundant contaminant sequences. First, we examined the fraction of reads from each set that aligned to a list of abundant murine ncRNAs. In each case we saw a clear reduction in the fraction of reads aligning to common ncRNAs **(Figure S1A)**. We then compared the fractions of reads that mapped uniquely to the genome for each sample before and after treatment depletion. As expected, the fraction of uniquely mapped reads was substantially increased for all samples, with more than 3-fold improvements in some samples **(Figure 2A)**. The fold change in uniquely mapping reads was far more pronounced than the changes in mapping to common ncRNAs; this is because even a small fold change in overall ncRNA percentage corresponds to a large fraction of the overall library being degraded **(Figure S1B).**

**Figure 2:**
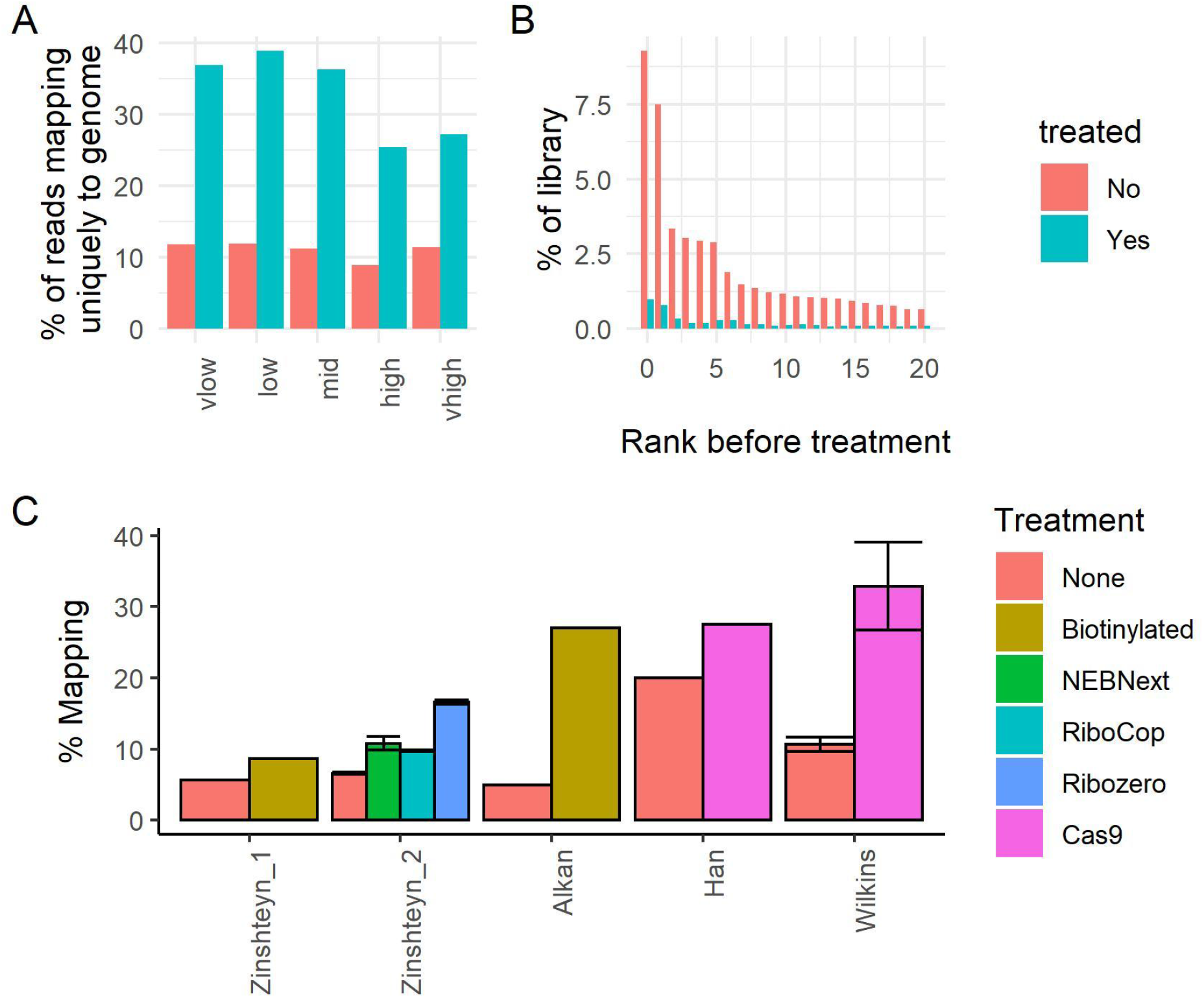
Ribocutter greatly increases the fraction of useful reads. **A:** The percentage of reads uniquely mapping to the genome (i.e. the percentage of useful reads) before and after treatment. **B:** The percentage abundance of the 20 most abundant contaminant sequences before and after treatment for a representative sample (medium RNase I). **C:** A comparison of different reported rRNA depletion efficiencies from different studies and methods. Error bars give standard deviation (where reported). “Wilkins” refers to the current study. “Biotinylated” refers to subtractive hybridisation using biotinylated probes. Note that the Zinshteyn experiments filter for reads within the CDS of genes (Zinshteyn et al. 2020), Alkan et al. filters for reads which align to protein coding transcripts (Alkan et al. 2021), and Han et al. (Han et al.2020) and Wilkins (this study) refer to all uniquely mapped reads to the genome which did not align to common ncRNA contaminants.

In our pilot experiments with lower concentration Cas9 and sgRNAs, we found that a significant fraction of our targeted sequences escaped degradation (data not shown). We therefore examined whether our higher-concentration protocol resolved this issue. We calculated the fraction of reads of the most abundant targeted sequences in our experiments before and after treatment. Pleasingly, without exception the most abundant targeted sequences were heavily depleted after treatment (**Figure 2B, Figure S2).** Overall, we found that our results compare favourably to other studies in terms of the increased fraction of useful reads after depletion (**Figure 2C**).

### Off-target depletion is minimal

Next, we analysed whether there was significant off-target degradation. Filtering for abundant sequences to minimise random noise (>0.01% prior to depletion), we compared the fraction of the library that each abundant sequence represented before and after treatment, both for targeted and non-targeted control sequences. Whereas targeted sequences were almost universally depleted after treatment (99%), the vast majority of non-targeted sequences had increased relative abundance after treatment (92%, **Figure 3A-B**). We found that ‘non-targets’ that were near-matches to sgRNAs were significantly more likely to be depleted (**Figure S3**, R = 0.48, p < 1e-15); it is likely that many of these ‘non-target’ reads are actually derived from genuine target rRNAs, but due to sequencing error appear as ‘non-target’ sequences. Notably, this experiment was performed with a large number of distinct sgRNAs (42 distinct sequences, required due to the different RNase I concentrations and tissues); in experiments with a smaller, more defined set of targeted sequences, the level of off-target degradation would presumably be lower.

**Figure 3:**
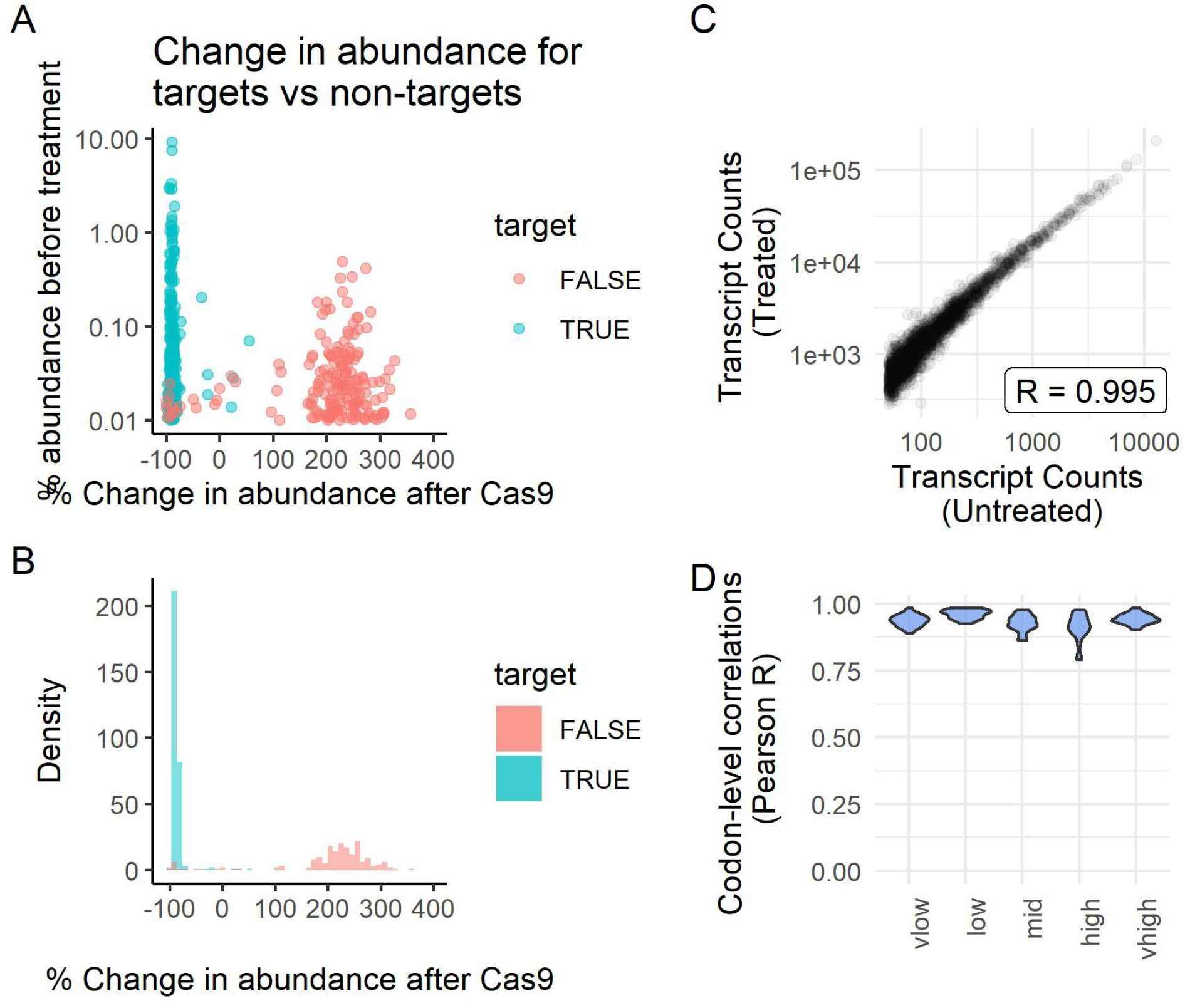
Ribocutter has minimal off-target effects. **A:** The average abundance of contaminant sequences (across both untreated and treated datasets) plotted against the average change in abundance after treatment for a representative sample (medium RNase I), stratified by whether they are targets of the sgRNA library. **B:** Histogram representation of A. **C:** The number of footprints aligning to each transcript, before (x axis) and after (y axis) treatment, for a representative sample (medium RNase I); Pearson’s R correlation shown. **D:** Codon-level correlations before and after treatment for the top 15 transcripts for each RNase I concentration (ranked by footprint density).

We examined whether the small extent of off-target degradation has any detectable impact on the quantification of ribosome profiling experiments. We analysed both transcript-level and codon-level correlations between the untreated and treated datasets. For transcripts-level counts, filtering for those with at least 50 footprints in each condition, we observed a very strong correlation before and after treatment (Pearson R > 0.99, Spearman Rho ~ 0.92-0.95; **Figure 3C, Figure S4**). Codon-wise correlations across each transcript are expected to be lower due to noise produced by the much decreased depth; we therefore filtered for the top 20 transcripts by footprint density for each sample. As expected, the codon-level correlations were somewhat weaker, however still proved to be highly reproducible before and after treatment (**Figure 3D).** Combined, these results demonstrate that the level of off-target degradation is very low compared to the on-target degradation, and has negligible impact on the quantitative information in data.

### Cas9 preserves footprint characteristics and periodicity

One method for depleting contaminating ncRNA from RNA-seq libraries is through RNase H treatment with antisense oligos that hybridise with specific contaminant sequences. However, it was recently reported that RNase H treatment is not suitable for Ribo-seq as it distorts the distribution of footprint sizes (and thus periodicity) in the library (Zinshteyn et al. 2020). We therefore examined whether Cas9 treatment produces similar distortions. We classified each read based on its sub-codon position relative to the start codon, its length, and whether it had a 5’mismatch (due to non-templated nucleotide addition during reverse transcription). We then plotted abundances of each footprint type across our five samples before and after treatment. We observed a near-perfect correlation (**Figure 4A**, R = 0.9997), demonstrating that Cas9 treatment does not distort the abundance of different footprint types.

**Figure 4:**
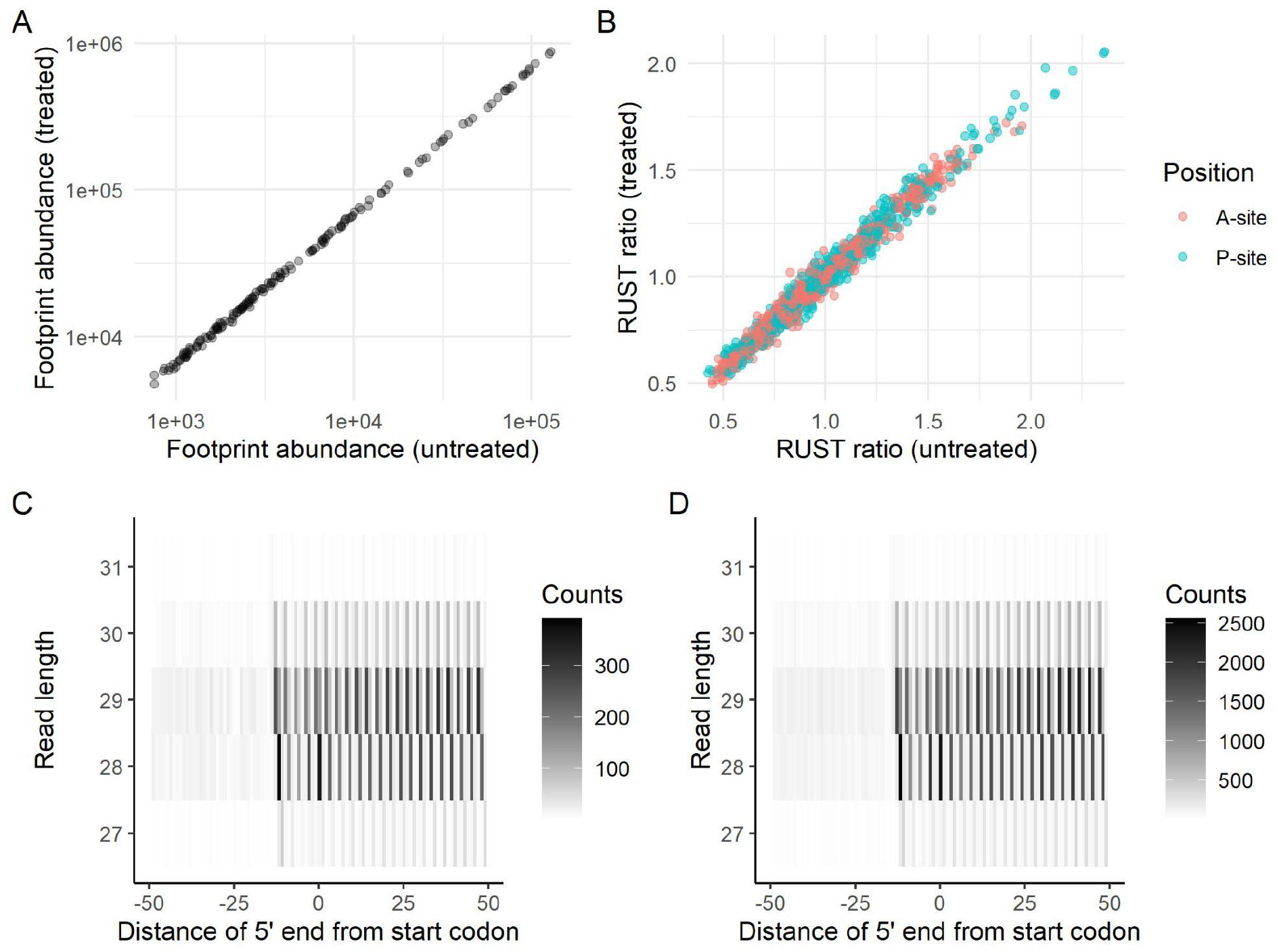
Ribocutter preserves key features of ribosome footprints. **A:** The abundance of specific footprint types before and after treatment for all five samples combined; each dot represents a single footprint type (eg. length, subcodon position, 5’ mismatch) from a single sample; R = 0.9997. **B:** Codon RUST enrichment ratios in the P and A sites of two abundant footprint types across all five samples; R = 0.988. **C** and **D:** heatmaps of footprint numbers around the start codons of transcripts for the “vhigh” sample (chosen due to its strongest periodicity), before and after Cas9 treatment respectively.

Next, we examined whether the codon biases present in the A and P sites of each footprint type, which correspond to positions that are slow to decode, correlated well between untreated and treated samples (O’Connor, Andreev, and Baranov 2016). Filtering for the major footprint types (28 nucleotides in frame 0 without a mismatch, and 29 nucleotides in frame 0 with a mismatch) we compared the P and A site codon RUST ratios, again finding a very strong correlation (**Figure 4B**, R = 0.988).

As a final quality control, we examined the patterning of footprints around the start codons of all protein coding transcripts, before and after treatment. Once again, we saw no evidence that Cas9 peturbs the sub-codon level resolution of the footprints (**Figure 4C-D**). Overall, these results suggest that Cas9-mediated depletion of contaminant ncRNAs is both highly effective and well suited to ribosome profiling.

## Discussion

### Cost per sample

The Ribocutter protocol can be rapidly and affordably performed on high numbers of samples as treatment uses a single multiplexed library as input. Here we assume that 24 samples are multiplexed, which is typical for a HiSeq or NovaSeq lane. Although there will be an up-front cost for some reagents, many of the reagents listed (for example Phusion, RNase A, DNA purification beads) are commonplace in research laboratories and will likely not have to be purchased specifically for this protocol. The net result is a dramatically less expensive protocol compared to commercial options: rRNA depletion from 24 samples using popular commercial kits would cost approximately £1,000, versus less than £12 with Ribocutter.

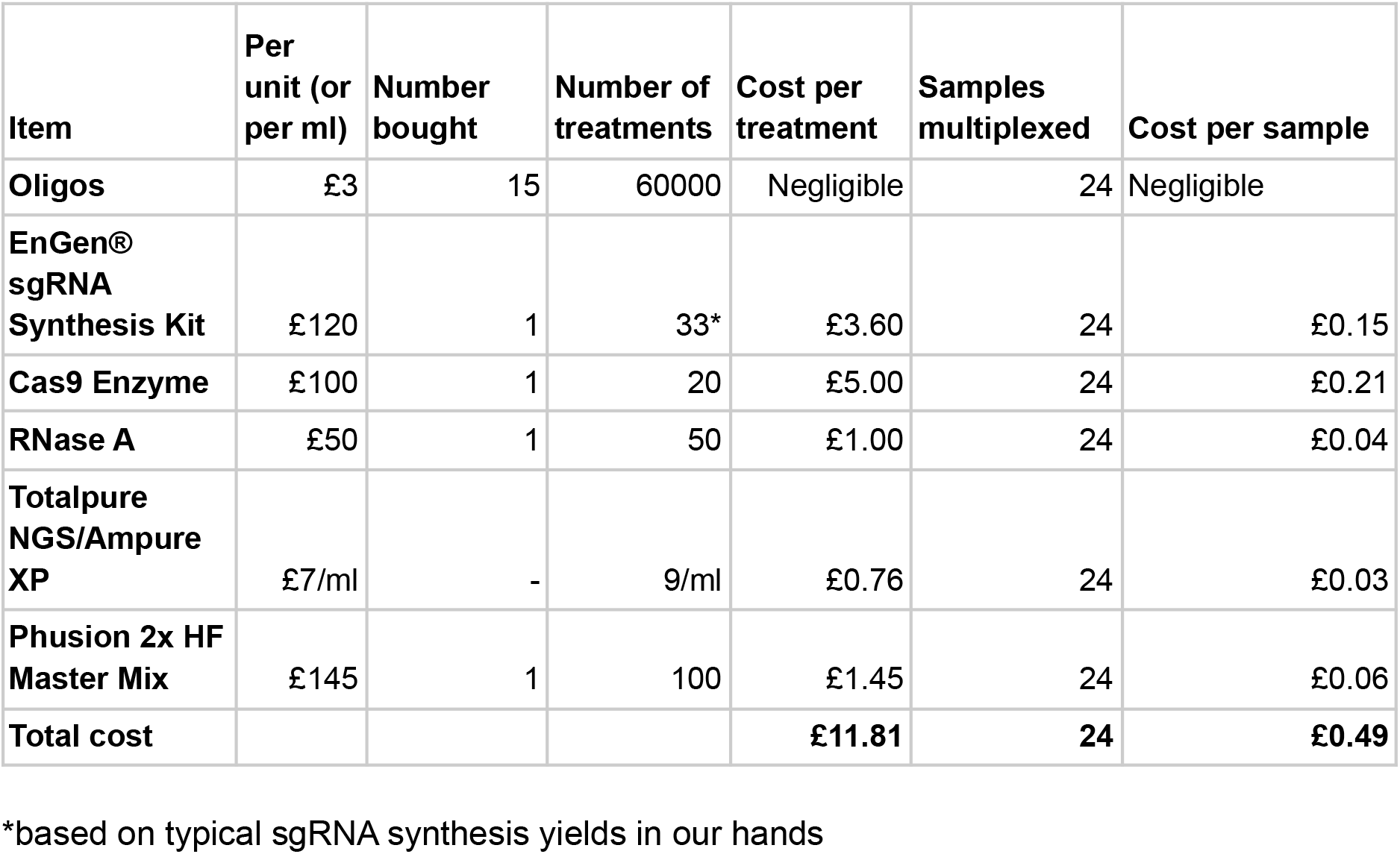

### Advances over previous protocols

The original study describing Cas9-mediated rRNA depletion for RNA-seq performed depletion on a non-amplified dsDNA library, prior to PCR purification (Gu et al. 2016). However, ribosome profiling libraries, and other similar small-RNA libraries such as iCLIP, are single-stranded prior to the PCR amplification step (König et al. 2010; McGlincy and Ingolia 2017). Given the limited activity of S. pyogenes Cas9 enzyme on ssDNA substrate, it is therefore necessary to amplify ribosome profiling libraries by PCR to produce dsDNA prior to Cas9 treatment (Ma et al. 2015;Hardigan et al. 2019). This has the added benefit that an aliquot of the amplified library, which will contain the full library complexity, can be held in reserve, reducing the risk of sample loss due to technical error.

In the Ribocutter protocol, we introduce two time-saving innovations. First, we recommend the use of a commercial 1-step sgRNA synthesis kit to produce sgRNAs - although this kit is designed for only a single template oligo, we have found that it is capable of producing a large pool of sgRNAs when using a pool of template oligos as input. This greatly reduces the time and cost associated with sgRNA synthesis.

Second, instead of a time-consuming gel purification step to enrich for uncleaved amplicons (Han et al. 2020), we recommend a quick PCR enrichment followed by beads purification. This also enables the DNA substrate concentration in the Cas9 reaction to be lower (as it will subsequently be amplified), resulting in a greater excess of sgRNAs and Cas9, which will improve on-target efficacy. A high ratio of sgRNA/Cas9 complex to substrate is important because S. pyogenes Cas9 is a single-turnover enzyme *in vitro* (Sternberg et al. 2014). This may explain why our depletion was more efficient than the previous publication using Cas9 with ribosome profiling, which used gel purification (Han et al. 2020), however various other experimental factors may have influenced this too. It is possible that a multiple-turnover Cas9 derivative might further improve degradation efficiency and/or lower reaction cost, though we have yet to explore this (Yourik et al. 2019).

### Cas9-mediated depletion compared to other depletion methods

Despite its high cost, Illumina Ribozero had been widely used by the ribosome profiling community, however it has been discontinued (McGlincy and Ingolia 2017; Zinshteyn et al.2020). Although similar products based on subtractive hybridisation, such as Lexogen Ribocop, are available commercially, they are often prohibitively expensive and, because they target the entire rRNA sequence rather than specific high-abundance contaminants, are not well suited to ribosome profiling due to risk of oversaturation (McGlincy and Ingolia 2017; Zinshteyn et al.2020; Alkan et al. 2021).

Many studies have recommended the use of biotinylated DNA oligos to perform targeted subtractive hybridisation, reducing cost and increasing on-target depletion levels (Ingolia et al.2012; McGlincy and Ingolia 2017; Zinshteyn et al. 2020; Alkan et al. 2021). Compared to Cas9, this approach may be somewhat faster overall, especially if pre-biotinylated oligos are purchased, and does not require the presence of a PAM motif in the targeted sequence. This may be particularly advantageous for depletion of ncRNAs which are too short to target with sgRNAs, as with Cas9 this would require the inclusion of flanking PAM-containing adapter sequences during sgRNA design (Hardigan et al. 2019).

Cas9, however, offers several clear advantages. It can be performed on a single multiplexed sample, eliminating technical variation, and its input is an aliquot of PCR-amplified DNA, reducing risk of sample loss due to technical error or RNA degradation. This also means that contaminants do not have to be predicted in advance, and particularly troublesome libraries, for example disome profiling libraries (Han et al. 2020), could be depleted of contaminants in multiple sequential rounds. Furthermore, Cas9 has very high specificity, meaning there is low risk of off-target depletion - indeed our data suggest that, even with a relatively large number of sgRNAs (>40), the level of off-target depletion is very low. One method of circumventing the requirement of a PAM motif within the targeted sequence itself is to include a PAM motif in the 5’ or 3’ adapter (Hardigan et al. 2019). However, we found that 90% of primary rRNA contaminants can be targeted by Cas9 without the need for additional PAM-containing adapter sequences, and therefore we did not use this approach in our study. Nevertheless, we do allow short 5’ and 3’ flanking sequences to be optionally included when designing sgRNA templates with the Ribocutter software (options --a5 and --a3), but, the code warns the user of a potential risk of increased off-target effects with this approach, and ensures that the added sequences are short.

Overall, we recommend Cas9 treatment as a viable and in many respects superior alternative to subtractive depletion with biotinylated oligos for ribosome profiling. We envisage that our streamlined protocol and easy-to-use software will enable this method to be more broadly adopted by the ribosome profiling community.

## Methods

### Generation of ribosome profiling libraries

Two snap-frozen mouse brains from P14 mice were lysed in ice-cold lysis buffer (McGlincy and Ingolia 2017), supplemented with 8% glycerol, using a glass homogeniser Ribosome footprints were generated essentially as described, using a sucrose cushion (McGlincy and Ingolia 2017). RNA concentration was measured using a QuantiFluor RNA system (Promega), then 0.1 (vlow), 0.5 (low), 0.75 (mid), 1.5 (high) or 3 (vhigh) ul of RNase I (EN0601, Thermo Fisher Scientific) per 30 ug of RNA were used for digestion. One mouse brain was used for vlow and low, and the second was used for mid, high and vhigh. Digestion was performed at 16 degrees for 45 min with gentle shaking, then quenched with SUPERase-In (Thermo Fisher Scientific).

Following RNA extraction from RNA pellets after ultracentrifugation, an Illumina-compatible sequencing library was produced following a protocol based on iCLIP/irCLIP (Blazquez et al.2018). Sequencing was performed at the Francis Crick Institute using an Illumina HiSeq 4000 machine (SR100).

### Raw data processing

Raw fastqs were demultiplexed and sequencing adaptors were trimmed using Ultraplex v1.1.5 (Wilkins et al. 2021). A Snakemake (Köster and Rahmann 2012) pipeline constituting of the following steps was used to align RFPs to the genome: reads were first aligned to a reference fasta of common ncRNA sequences using Bowtie 2 v2.4.2 (Langmead and Salzberg 2012), then reads which escaped pre-mapping were aligned to the mouse GRCm38 genome assembly using STAR v2.6.1a (Dobin et al. 2013). Next, aligned reads were deduplicated using UMI-Tools v1.0.1 (Smith, Heger, and Sudbery 2017), and final quality control was performed using in house scripts (Riboloco).

### Cas9 rRNA depletion

sgRNA templates were designed using the Ribocutter software and synthesised using the EnGen sgRNA synthesis kit, using an extended 2 hour incubation to maximise yield (Table S1). rRNA depletion was performed as described above, using a 33 nM PCR product as input. The optional murine RNase inhibitor (described above) was included in all relevant steps. Following reamplification with P5/P3 primers and purification (described above), the treated library was resubmitted for sequencing using the same machine/settings.

~~~
P3 primer: 5′-CAAGCAGAAGACGGCATACGAGATCGGTCTCGGCATTCCTGCTGAACCGCTCTTCCGATCT-3′
P5 primer: 5′-AATGATACGGCGACCACCGAGATCTACACTCTTTCCCTACACGACGCTCTTCCGATCT-3′
~~~

### Data analysis

Sequence copy numbers were analysed by the Ribocutter software (this study; v0.0.8) using options “--save_stats --stats_frac 0.000001”. Alignment statistics were calculated by analysing the output logs of Bowtie2, STAR and UMI-Tools. All additional analysis was performed using a custom R script.

## Data availability

Adaptor-/quality-trimmed demultiplexed fastq files are available at E-MTAB-10736.

## Author contributions

OGW designed the study and performed all experimentation and data analysis. JU supervised the study. OGW and JU wrote the manuscript.

## Acknowledgements

We thank Pietro Fratta and Rafaela Fernandez de la Fuente for providing the mouse brain samples.

---

**Supplementary Figure 1:**
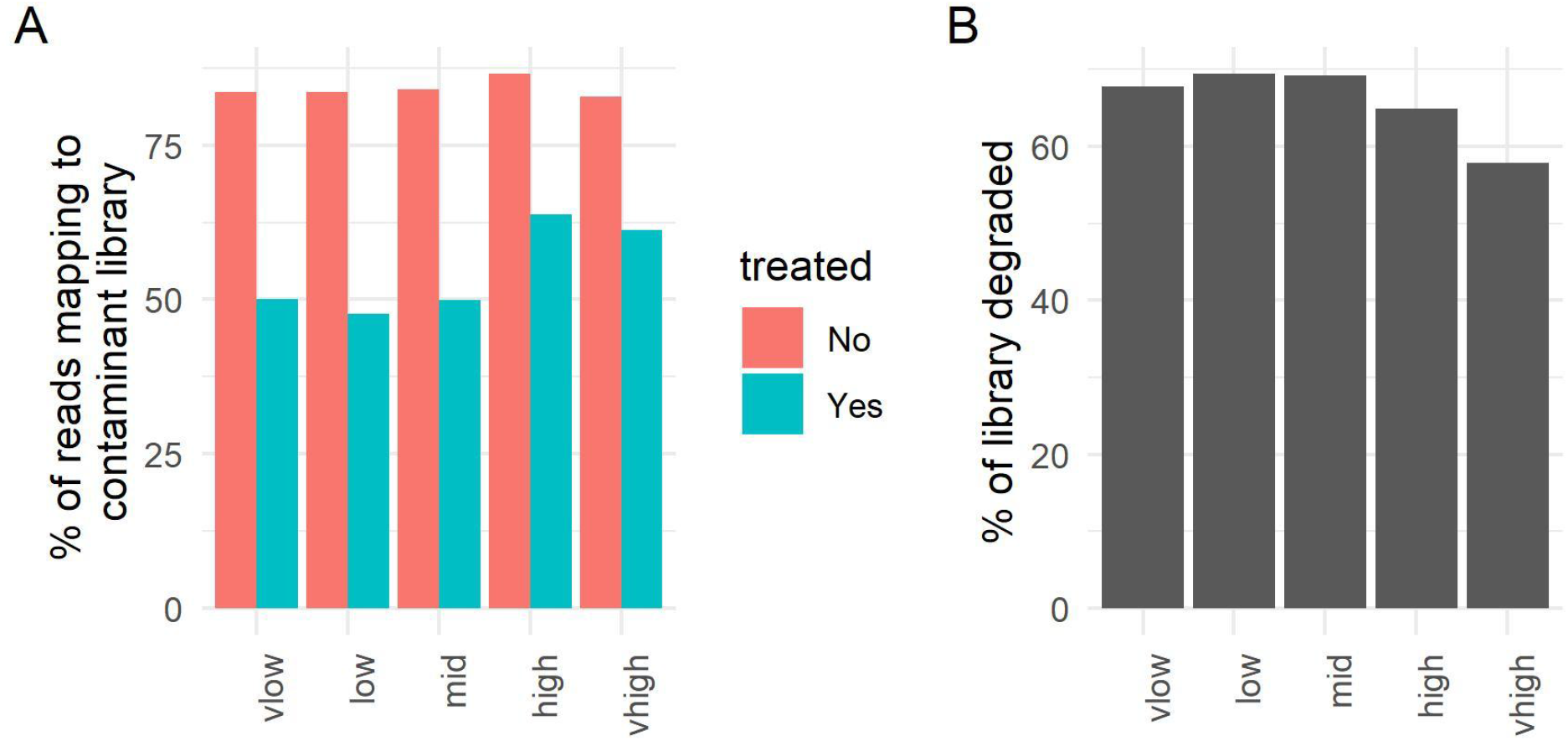
Statistics on efficiency of Cas9 degradation. **A:** The percentage of reads mapping to a library of common contaminant ncRNAs before and after treatment. **B:** An estimate of the fraction of each library that was degraded by Cas9 treatment.

**Supplementary Figure 2:**
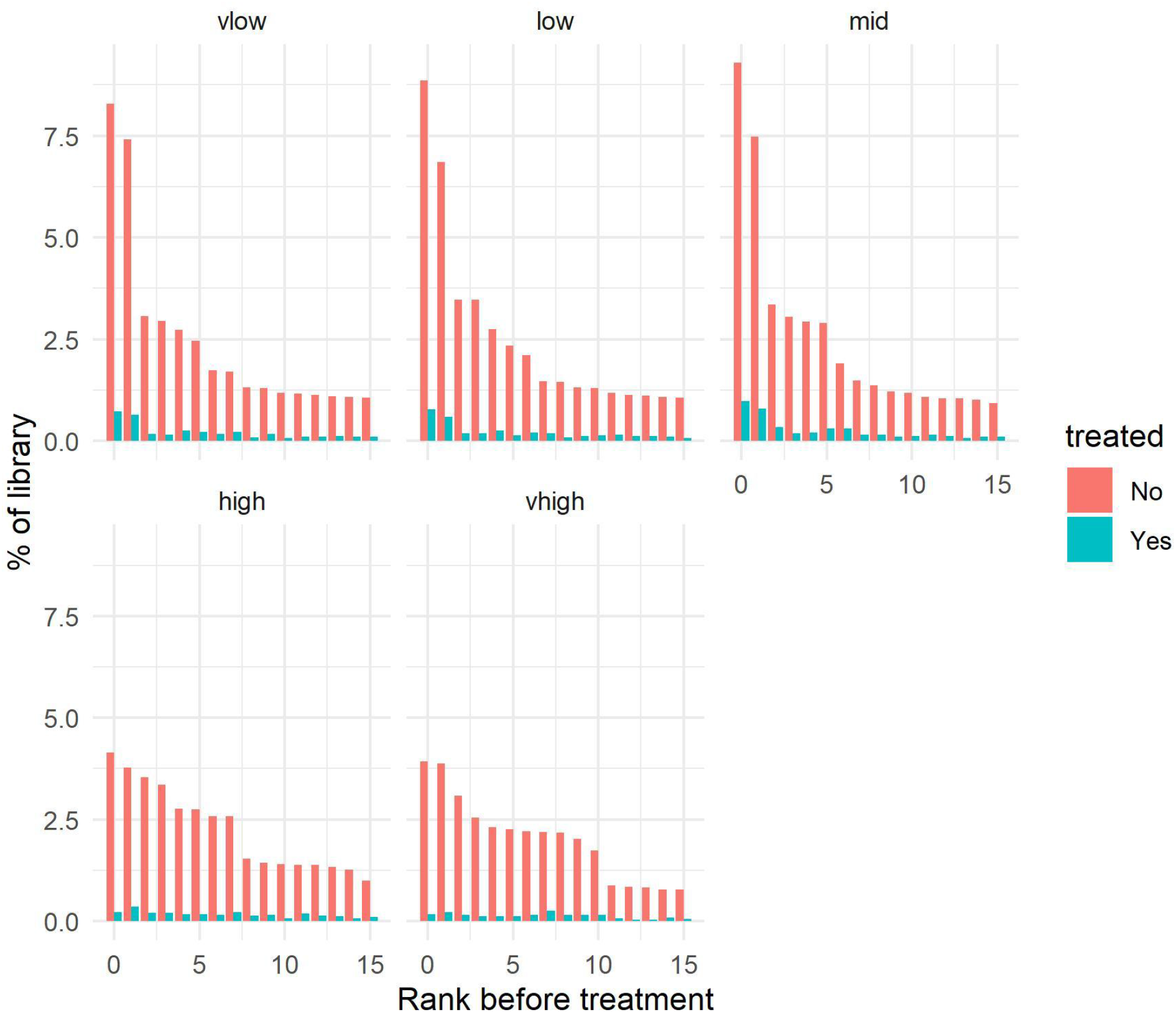
Levels of most abundant contaminant sequences before and after treatment. Equivalent of Figure 2B but with all five samples.

**Supplementary Figure 3:**
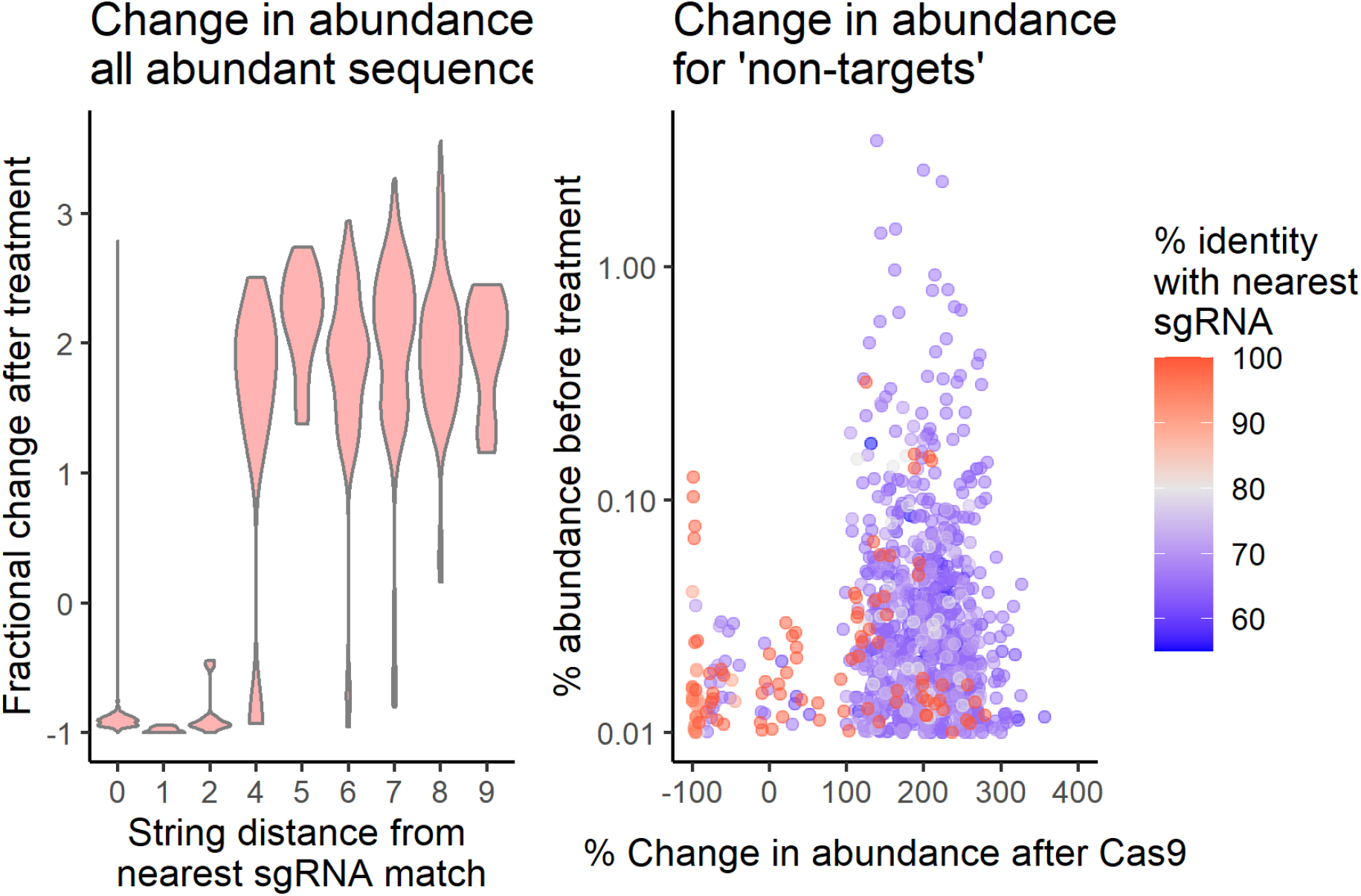
Near-targets are more likely to be depleted. Left: violin plot of all abundant sequences (>0.01% prior to depletion), separated by their string distance from the closest sgRNA match, and their respective change in abundance after treatment. Right: scatter graph showing all abundant ‘non-target’ sequences’ (>0.01% prior to depletion) abundance against change in fractional abundance, coloured by their percentage identity to the nearest sgRNA. *Note that apparent ‘non-target’ sequences may be genuine targets that were incorrectly sequenced*.

**Supplementary Figure 4:**
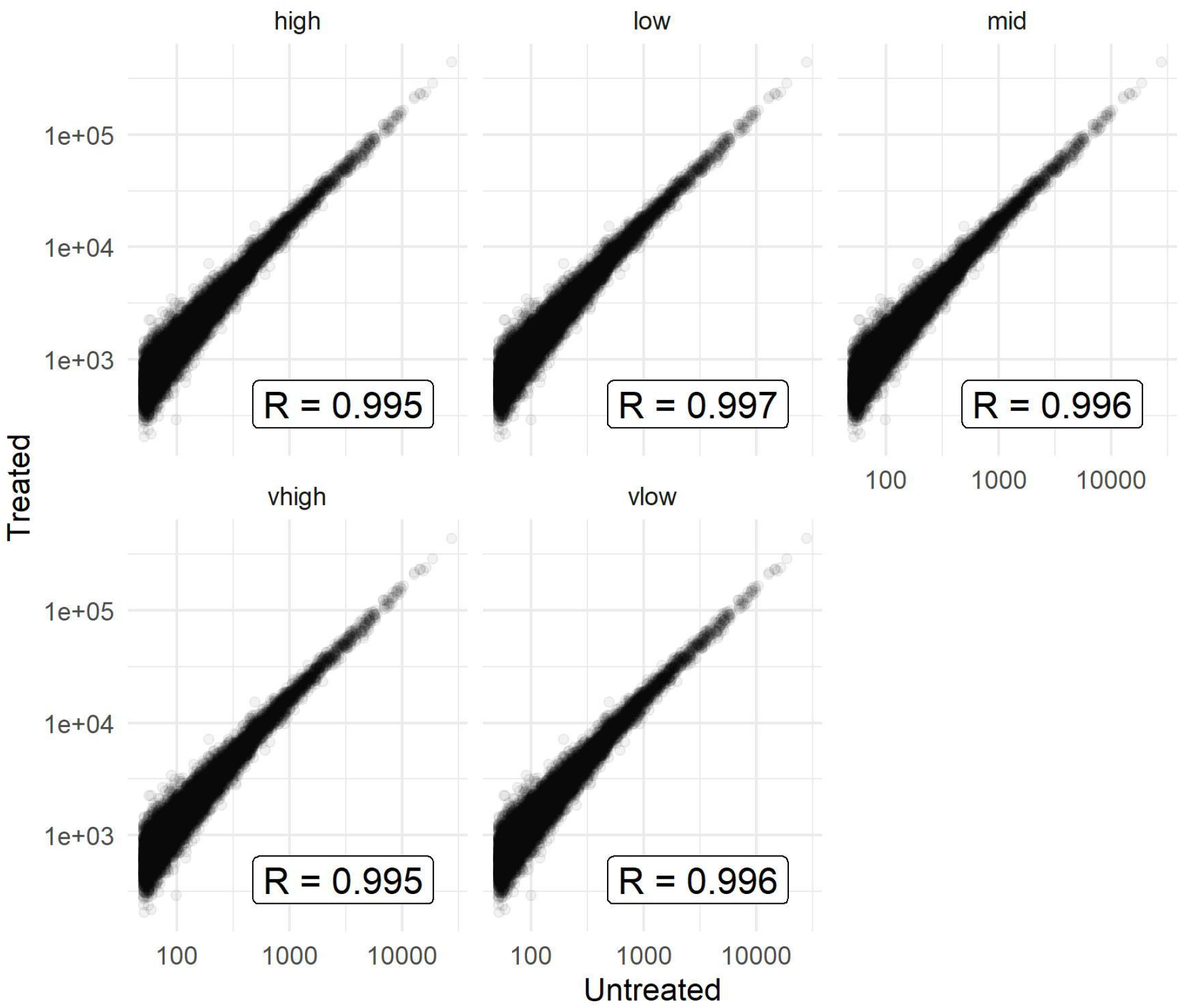
Transcript-level footprint counts before and after treatment. Equivalent of Figure 3C but with all five samples.

**Table S1.**
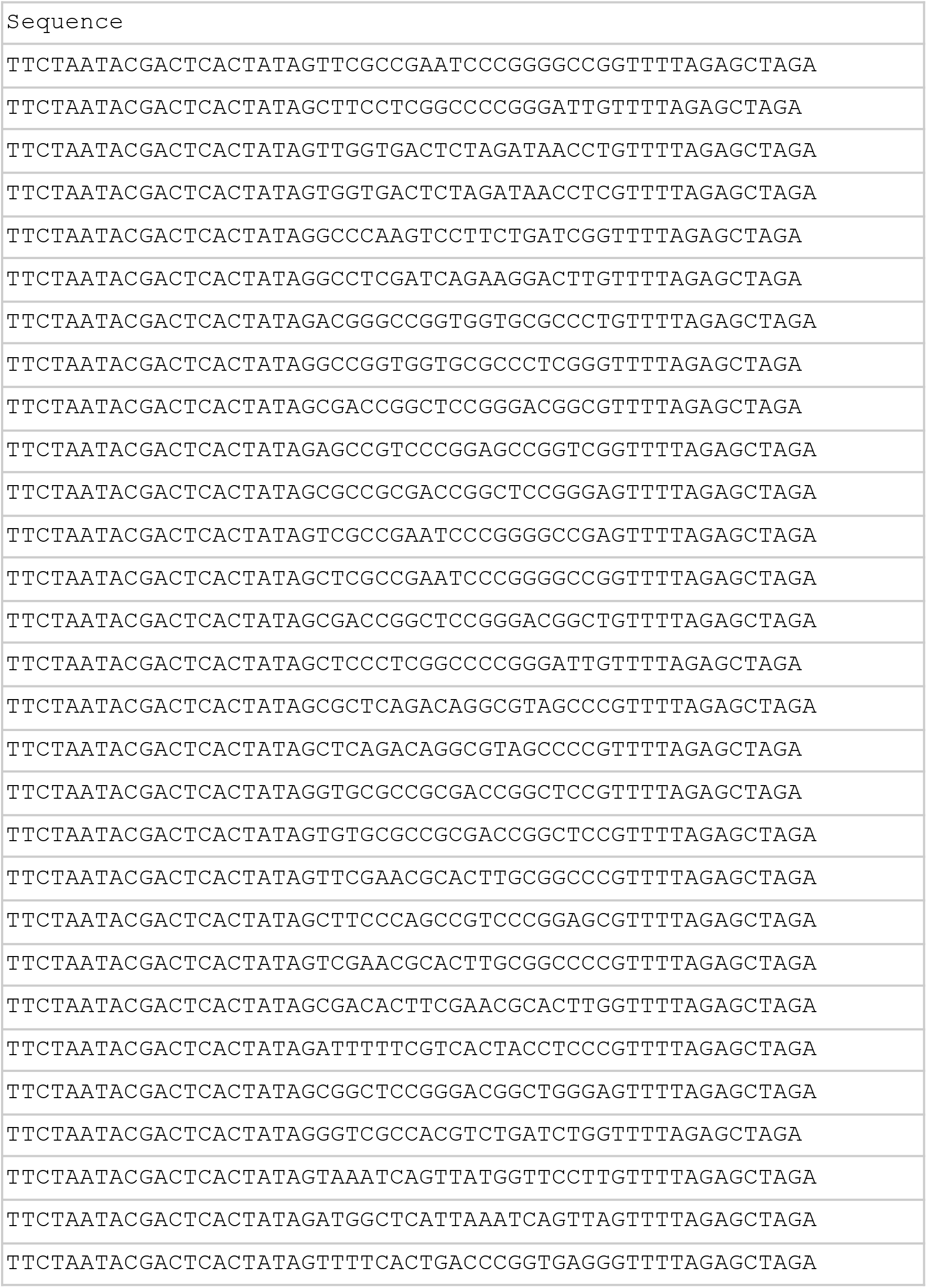
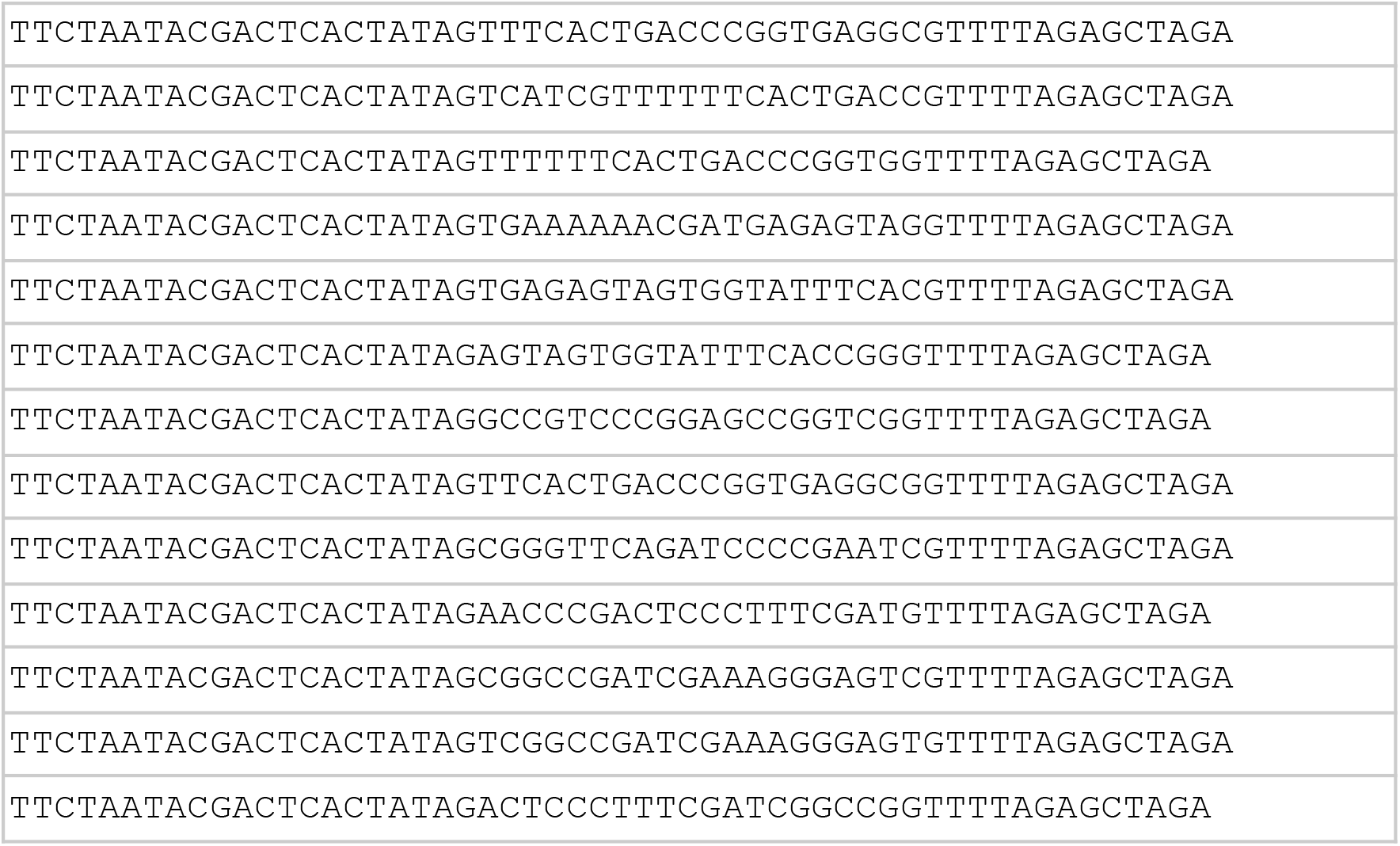
Oligos for the synthesis of sgRNAs targeting common mammalian ncRNA sequences.

## Footnotes

* *Note that each synthesis reaction generates enough sgRNA for many treatments - it is not necessary to produce fresh sgRNA for each treatment*.

** In some cases it may be possible to avoid PCR amplification and instead perform a gel extraction of the uncleaved DNA, then proceed directly to sequencing submission.

